# SIN3 gene regulatory activity is linked to RNA polymerase II pausing

**DOI:** 10.1101/2023.02.03.526989

**Authors:** Imad Soukar, Anindita Mitra, Lori Pile

**Affiliations:** Department of Biological Sciences, Wayne State University, Detroit, Michigan, 48202, USA

**Keywords:** SIN3 isoforms, Epigenetic modifications, RNA polymerase II, Pausing index, Soft regulation

## Abstract

The chromatin environment has a significant impact on gene expression. Chromatin structure is highly regulated by histone modifications and RNA polymerase II binding dynamics. The SIN3 histone modifying complex regulates the chromatin environment leading to changes in gene expression. In *Drosophila melanogaster*, the *Sin3A* gene is alternatively spliced to produce different protein isoforms, two of which include SIN3 220 and SIN3 187. Both SIN3 isoforms are scaffolding proteins that interact with several other factors to regulate the chromatin landscape. The mechanism through which the SIN3 isoforms regulate chromatin is not well understood. Here, we analyze publicly available data sets to allow us to ask specific questions on how SIN3 isoforms regulate chromatin and gene activity. We determined that genes repressed by the SIN3 isoforms exhibited enrichment in histone H3K4me2 and H3K4me3 near the transcription start site. We observed an increase in the amount of paused RNA polymerase II on the promoter of genes repressed by the isoforms as compared to genes that require SIN3 for maximum activation. Furthermore, we analyzed a subset of genes regulated by SIN3 187 that suggest a mechanism in which SIN3 187 might exhibit hard regulation as well as soft regulation. Data presented here expand our knowledge of how the SIN3 isoforms regulate the chromatin environment and RNA polymerase II binding dynamics.

**Summary statement:** SIN3 cofactors can activate or repress genes. Using bioinformatic analysis, we find that histone methylation and RNA polymerase II binding profiles differ at SIN3-regulated genes with distinct transcriptional outcomes.

## INTRODUCTION

Nucleosomes are composed of DNA strands that wrap around histone proteins. This interaction facilitates DNA compaction into the nucleus of the cell. Histone proteins are subject to post-translational modifications such as acetylation and methylation. The presence or absence of specific histone modifications can impact gene activity (Tse et al., 1998). The regulation of gene activity by histone modifications occurs through two major mechanisms. First, the modification of histone N-terminal amino acids (histone tails), such as the acetylation of lysine residues, brings about the nullification of the ionic interaction between the positively charged lysine and the negatively charged DNA. The loss of such interaction leads to a less compact nucleosome, making the DNA more accessible for RNA polymerase binding (Nightingale et al., 1998). The second mechanism is through signaling and transcription regulator recruitment. Histone tail modifications can act as a signal that recruits transcription factors and chromatin effectors to a gene locus, conferring an effect on gene regulation (Cheung et al., 2000).

One such chromatin effector complex is the SIN3 histone deacetylase (HDAC) complex, which regulates the acetylation and methylation of histone tails (Silverstein and Ekwall, 2005). The SIN3 complex is composed of the SIN3 scaffolding protein, which recruits other proteins, including histone deacetylase 1 (HDAC1), and the histone demethylase dKDM5A/LID (Moshkin et al., 2009; Spain et al., 2010). *Sin3A* is an essential gene in metazoans. The knockout of *Sin3A* leads to loss of viability in both Drosophila and mouse models (Dannenberg et al., 2005; Neufeld et al., 1998; Pennetta and Pauli, 1998). Drosophila SIN3 regulates genes encoding proteins in many important pathways including cell cycle, one carbon, and central carbon metabolism (Pile et al., 2003). Consistent with the changes in gene expression, RNA interference-mediated reduction of *Sin3A* in Drosophila S2 cultured cells leads to a G2 arrest in the cell cycle (Pile et al., 2002). Furthermore, reduction of *Sin3A* in S2 cells leads to the deregulation of many one carbon and central carbon metabolites, indicating a role for the SIN3 complex in metabolic regulation of the cell (Liu and Pile, 2017; Liu et al., 2020).

The Drosophila *Sin3A* gene encodes multiple isoforms of SIN3 through differential splicing (Pennetta and Pauli, 1998). Two of the most prevalent isoforms are SIN3 220 and SIN3 187, named based on the predicted molecular weight. The isoforms have differential expression patterns and distinct abilities to rescue a genetic mutation in *Sin3A.* SIN3 isoforms are expressed at similar levels in the initial stages of Drosophila embryogenesis (Sharma et al., 2008). SIN3 220 levels increase in stages 12-16 but fall at the final stage of embryogenesis, at which time, the expression of SIN3 187 becomes predominant (Sharma et al., 2008). The lethality due to genetic disruption of the *Sin3A* gene can be rescued by a transgene designed to express SIN3 220, while a transgene encoding SIN3 187 is unable to suppress the lethal phenotype (Spain et al., 2010). Additionally, while both isoforms interact with HDAC1 along with other core SIN3 complex components, our group has shown that these isoforms bind unique proteins as well. For example, dKDM5A/LID is found in the SIN3 220 complex and not the SIN3 187 complex (Spain et al., 2010).

The SIN3 isoforms are recruited to thousands of genes throughout the Drosophila genome, many of which are common targets between the two isoforms, suggesting an overlapping mechanism of regulation by the isoforms (Saha et al., 2016). Specific gene targets that are differentially regulated by the isoforms and belong to distinct gene ontology (GO) categories, however, have also been identified. The differential interaction of specific proteins with the isoforms could lead to the differential regulation of the observed GO pathways (Saha et al., 2016). Both of the SIN3 isoforms have been implicated in gene repression and gene activation, yet the chromatin context in which SIN3 complexes can activate or repress genes is not well understood. RNA polymerase II (RNA Pol II) transcribes eukaryotic genes in three general steps. First, RNA Poll II is recruited to gene promoters to form the pre-initiation complex (PIC) where the RNA polymerase interacts with DNA at the transcription start site (TSS). Next, the C-terminal domain (CTD) of the polymerase is phosphorylated at serine (Ser) 5, transcription is initiated and then RNA Pol II pauses 20-60 base pairs downstream of the TSS. This promoter proximal pausing is dependent on pausing factors such as NELF-A and the DSIF complex containing SPT5 (Adelman and Lis, 2012). To release the polymerase from its paused state, the transcription elongation factor TEFb phosphorylates RNA Pol II, NELF-A and the DSIF complex, leading to the disassociation of NELF-A. Once transcription is terminated, RNA Pol II is released and recycled for another round of transcription. It has been long suggested that histone acetylation affects RNA Pol II dynamics, and many publications support this idea (Nightingale et al., 1998; Vaid et al., 2020). Studies from Nightingale et al. (1998) showed that histone acetylation positively affects RNA Pol II initiation. Furthermore, results of a recent study suggest a link between histone deacetylation activity and RNA Pol II pausing on a subset of developmental and signaling genes (Vaid et al., 2020). The authors found that HDAC inhibition leads to the release of RNA Poll II from the promoter paused state, leading to the elongation of select gene transcripts.

SIN3 is well-studied in its role as a gene regulator and scaffolding protein. However, the way in which SIN3 isoforms repress or activate genes has not been fully elucidated. We recently explored a potential mechanism of gene regulation by SIN3 (Mitra et al., 2021). We predicted that histone modifying complexes such as the SIN3 complex regulate housekeeping genes in a soft repression manner rather than an on/off switch. Soft repression is a mechanism whereby gene regulators affect the expression of genes by a small fold change to fine-tune the transcriptional output (<2 log2 fold change). For that study, we focused on genes repressed by the SIN3 220 isoform. Here, we investigate the mechanism of differential regulatory activity of the SIN3 isoforms by metagene analysis of SIN3-regulated genes. Our results support the idea that histone modifications regulated by SIN3 impact RNA Pol II dynamics. Additionally, our findings are consistent with our prediction that SIN3 acts as a soft repressor. Interestingly, we also obtain data to suggest that the SIN3 187 isoform acts as a hard regulator on some genes. In summary, we provide evidence to suggest that SIN3 fine tunes the expression of genes by regulating histone methylation and elongation dynamics of RNA Pol II.

## RESULTS

### SIN3 220 and SIN3 187 binding is enriched at the TSS of repressed and activated gene targets

To understand how SIN3 isoforms can lead to either gene repression or gene activation, we sought to uncover possible differences in protein binding and the chromatin landscape comparing SIN3 repressed and SIN3 activated direct gene targets. We did this by analyzing SIN3 isoform binding to genomic loci and comparing the common and unique targets. In this study, we expanded on our published analysis of the binding of the SIN3 isoforms by including targets unique to one isoform. In our previous study, we determined that the majority of SIN3 220 targets are also regulated by SIN3 187, while SIN3 187 regulates several genes that are specific to that isoform (Saha et al., 2016). To analyze the binding profile of the SIN3 isoforms, published genomic data were downloaded from NCB?s GEO database (Fig. 1A) and analyzed using the Galaxy platform (Afgan et al., 2018). To determine the list of genes directly regulated by the SIN3 isoforms, we integrated SIN3 RNA-seq data (Gajan et al., 2016) with ChlP-seq data (Saha et al., 2016). SIN3 220 directly regulates 405 genes; 60% (242/405) of those genes are repressed by SIN3 220, while 40% (163/405) are activated. Furthermore, 859 genes are directly regulated by SIN3 187, 55% (469/859) are repressed, while 45% (390/859) are activated. Of the 405 genes directly regulated by SIN3 220, 83% (335/405) are also regulated by SIN3 187 with only 17% (70/405) of genes uniquely regulated by SIN3 220 (Fig. 1B). For SIN3 187 targets, only 39% (335/859) of genes are also regulated by SIN3 200, with the majority of genes, 61% (524/859), uniquely regulated by SIN3 187.

**Fig. 1.**
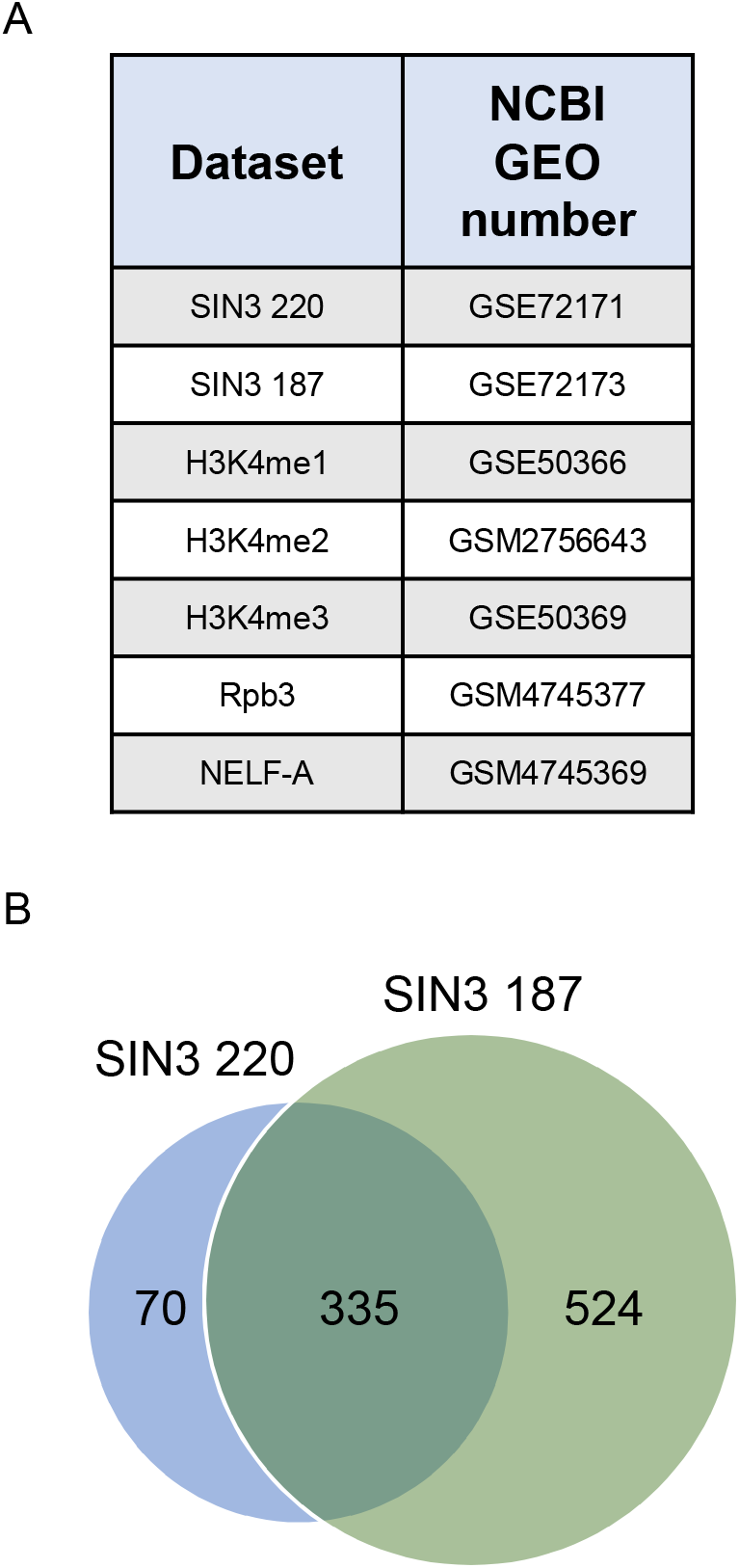
Datasets used in this study. A. Data sets used in this study are shown with the associated NCBI GEO number. B. Genes bound and regulated by SIN3 220 and SIN3 187 are plotted in a Venn diagram.

Our previous binding analysis showed that SIN3 isoforms are enriched at the TSS of genes (Saha et al., 2016). To follow up from that study, we asked, what are the differences between the enrichment of SIN3 at genes regulated by both isoforms, and genes regulated by SIN3 187 alone? Consistent with our previous analysis, SIN3 220 binding is enriched at the TSS of repressed genes and activated genes (Fig. 2A, B) compared to the gene body. Additionally, SIN3 220 binding is more enriched at repressed genes compared to activated genes (Fig. 2B). Likewise, SIN3 187 binding is more enriched at the TSS for both repressed and activated genes (Fig. 2A, 2B) compared to the gene body. To a lesser extent but similar to the SIN3 220 binding profile, genes repressed by SIN3 187 exhibited more binding at the TSS compared to activated genes (Fig. 2B). Next, we looked at genes exclusively regulated by SIN3 187 and not by SIN3 220. These genes are bound by SIN3 187 and change in expression only when SIN3 187 levels are perturbed and not when SIN3 220 levels are reduced (Saha et al. 2016). Interestingly, at this set of targets we do not observe a notable difference in SIN3 187 binding comparing the levels at repressed to activated genes (Fig. 2A, 2B). These data indicate that for the common repressed SIN3 gene targets, SIN3 complex activity might be impacting the PIC or other factors at the TSS. At activated targets and at the genes specific to SIN3 187, the location of gene regulatory activity may be more variable as complex binding is less TSS directed.

**Fig. 2.**
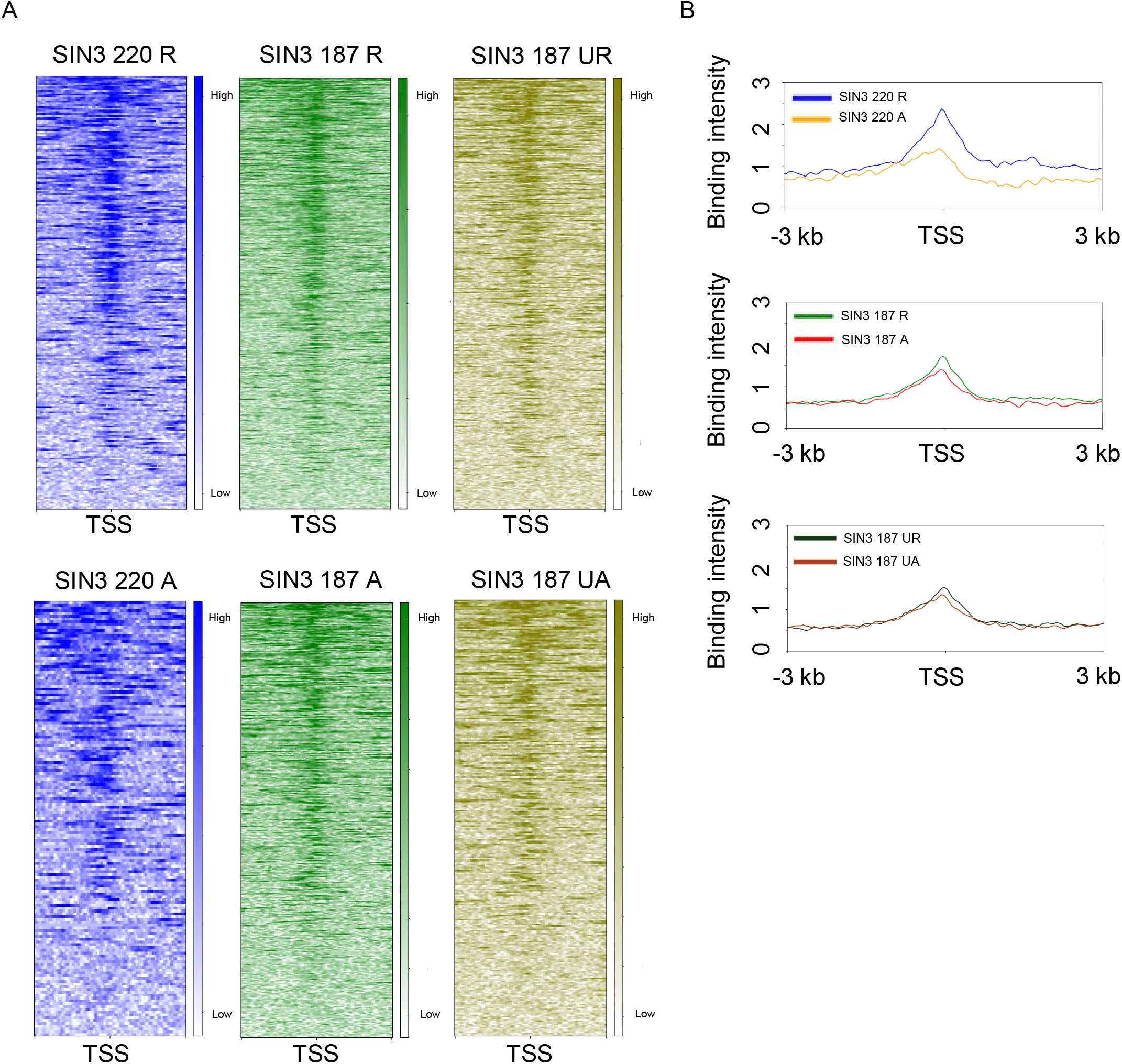
SIN3 isoform binding is different between activated and repressed genes. A. SIN3 220 and SIN3 187 binding on genes that change in expression when SIN3 220 and SIN3 187 levels are perturbed, visualized using heat maps. Uniquely regulated genes are those that are bound by SIN3 187 and only change in expression when SIN3 187 levels are changed. B. Average binding intensity of SIN3 isoforms on regulated genes plotted from −3 kb to 3 kb. R = repressed, A = activated, UR = uniquely repressed, UA = uniquely activated, TSS = transcription start site.

### SIN3 repressed genes have more enrichment of H3K4me2 and H3K4me3 compared to activated genes

SIN3 is a scaffold protein for assembly of a histone modifying complex that binds to gene promoters and can affect the neighboring chromatin environment (Gajan et al., 2016; Liu and Pile, 2017). SIN3 is associated with two proteins shown to regulate the post-translational modification profile of histone proteins, histone deacetylase HDAC1 and histone demethylase dKDM5/LID. Here, we wished to analyze the promoter histone modification profiles at SIN3-regulated genes. Due to lack of histone acetylation data generated from S2 cells, we were not able to examine the acetylation profile of SIN3 gene targets. We previously determined that the histone demethylase dKDM5/LID is a part of the SIN3 220 complex but not SIN3 187 complex (Spain et al., 2010). This finding led us to ask if genes regulated by the SIN3 isoforms have differential enrichment of methylation at target genes. To address this question, we determined the level of histone methylation at the direct SIN3 isoform targets. We chose H3K4me1, H3K4me2, and H3K4me3 to gain a full understanding of the H3K4 methylation pattern of SIN3-regulated genes. We expected genes to have low to no H3K4me1 levels, since this mark has been associated with gene enhancers rather than at the TSS (Heintzman et al., 2007). Additionally, we expected genes to have higher enrichment of H3K4me2 and H3K4me3 when compared to H3K4me1 levels. Our reasoning is that both of these marks are associated with actively expressed genes (Bernstein et al., 2005), and SIN3 has been shown to fine-tune the expression of expressed genes (Mitra et al., 2021). We downloaded H3K4 methylation data from NCBI GEO (Fig. 1A) and overlapped it with genes bound by SIN3 to generate heat maps and binding profiles. We found no difference in H3K4me1 enrichment between activated and repressed genes (Fig. 3A).

**Fig. 3.**
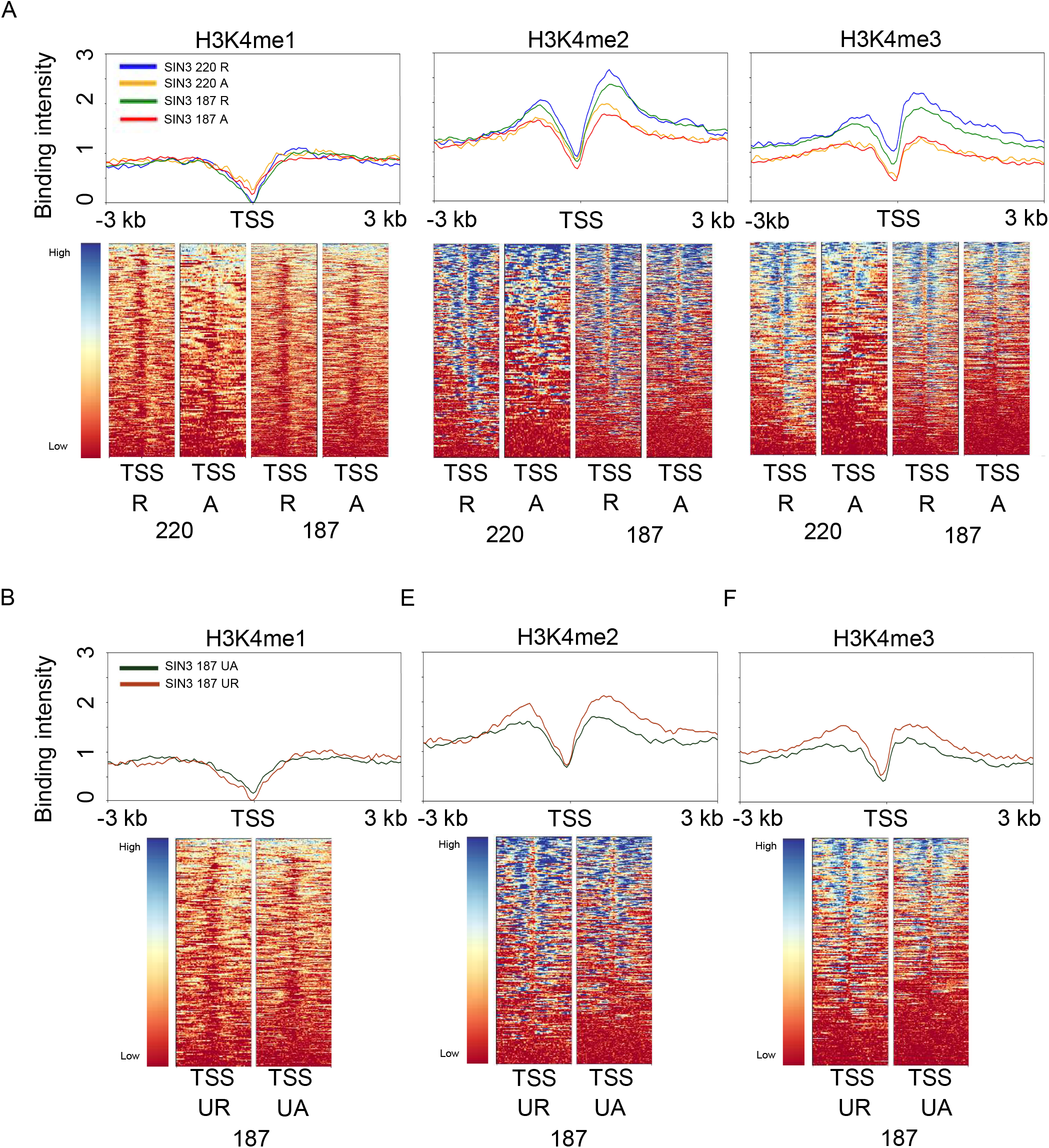
H3K4 methylation at genes regulated by the SIN3 isoforms. A. H3K4me1, H3K4me2, and H3K4me3 enrichment was mapped to SIN3 220 and SIN3 187 regulated genes. Heat maps and binding profiles were generated spanning genes from −3 kb to 3 kb. B. H3K4me1, H3K4me2, and H3K4me3 enrichment was mapped to genes bound by SIN3 187 and that only change in expression when SIN3 187 levels are changed. Heat maps and binding profiles were generated spanning genes from −3 kb to 3 kb. R = repressed, A = activated, UR = uniquely repressed, UA = uniquely activated, TSS = transcription start site.

On the other hand, we found higher enrichment of H3K4me2 and H3K4me3 at genes repressed by SIN3 220 compared to those genes that are activated (Fig. 3A). This enrichment was seen ~700 bp downstream of the TSS. For SIN3 187, we observe a similar trend wherein H3K4me1 enrichment was similar at repressed and activated targets, while SIN3 187 repressed genes showed higher enrichment of H3K4me2 and H3K4me3, compared to genes activated by SIN3 187 (Fig. 3A). To further dissect the histone methylation profiles of SIN3-regulated genes, we looked at genes exclusively regulated by SIN3 187 and not SIN3 220. To our surprise, we saw a similar but smaller trend in which H3K4me2 and H3K4me3 were enriched at repressed genes and not activated genes (Fig. 3B), though the level of these marks at the unique targets was lower in comparison to the common targets. Because SIN3 187 was not found to interact with dKDM5/LID, these results strongly suggest that dKDM5/LID is not the sole factor that affects the histone methylation profile of SIN3-regulated genes.

### RNA Pol II and negative elongation factor (NELF) are paused at the TSS of SIN3 220 and SIN3 187 repressed genes and not at activated genes

Transcription elongation by RNA pol II is regulated in the cell by pausing factors such as negative elongation factor-A (NELF-A) and DRB sensitivity-inducing factor (DSIF) (Adelman and Lis, 2012). HDAC1 influences the release of RNA pol II from the TSS of a subset of development and signaling genes (Vaid et al., 2020). Since HDAC1 is one of the core components of both SIN3 220 and SIN3 187 complexes, we asked if RNA pol II is paused at SIN3-regulated genes. To do this, previously published RNA Pol II and NELF-A genome-wide binding data (Mazina et al., 2021) was downloaded and overlapped with the SIN3 binding data (Fig. 1 A). We observed an enrichment of RNA Pol II 50 bp downstream of the TSS of genes repressed by SIN3 220 (Fig. 4A). RNA Pol II was also enriched 50 bp downstream of the TSS of genes repressed by SIN3 187 (Fig. 4A). To quantify the difference in enrichment of RNA Pol II at SIN3 220 and SIN3 187 regulated genes, we determined the pausing index using a method modified from Vaid et al. (2020). To calculate the pausing index, the binding enrichment value at promoter region (−50 bp to +50 bp) was divided by the enrichment value along the genic region (+300 bp to −100 TES). SIN3 220 and SIN3 187 repressed genes exhibited higher RNA Pol II pausing when compared to genes that were activated (Fig. 4B). These data suggest that SIN3 repression of genes involves the regulation of RNA Pol II elongation dynamics.

**Fig. 4.**
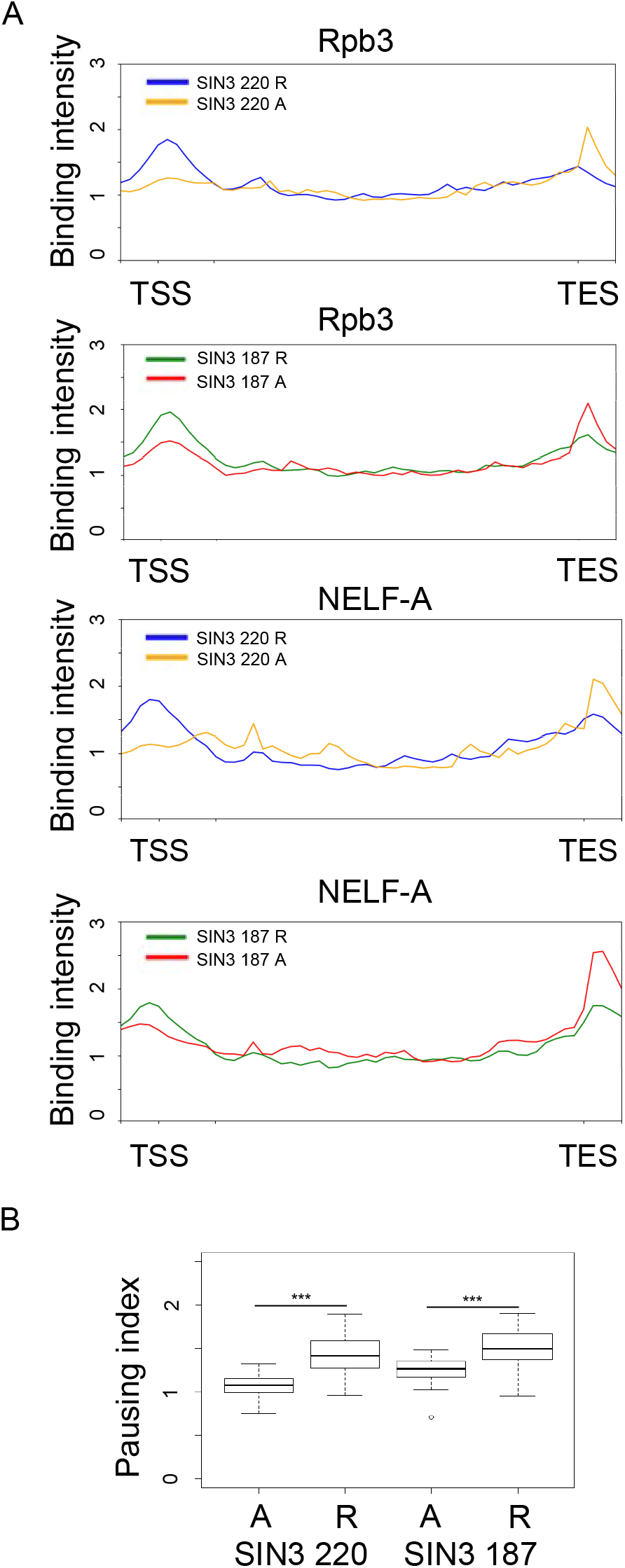
RNA pol II pausing at SIN3 regulated genes. A. RNA Pol II subunit Rpb3 and pausing factor NELF-A binding were mapped against genes regulated by both SIN3 isoforms. B. Pausing index of genes regulated by both isoforms. Rpb3 binding enrichment value at the promoter (−50 bp to +50 bp) was divided by Rpb3 enrichment value at the gene body (+300 bp to −100 bp TES). R = repressed, A = activated, TSS = transcription start site, TES = transcription end site. Significance was determined by Mann Whitney U Test. *** indicates p value <0.001.

To further investigate whether SIN3 repressed genes exhibit high RNA Pol II pausing, we asked if the binding of the pausing factor NELF-A is also different between repressed and activated genes. NELF-A is a pausing factor that plays an important role in RNA Pol II promoter proximal pausing (Yamaguchi et al., 2013). In line with our expectations, the NELF-A binding pattern was similar to that seen with RNA Pol II. NELF-A had a higher enrichment near the TSS of SIN3 220 and SIN3 187 repressed genes compared activated genes (Fig. 4A). These data support the idea that SIN3 220 and SIN3 187 repress some genes through a similar mechanism. Additionally, these findings suggest a mechanism in which SIN3 represses genes by regulating RNA Pol II pausing, possibly involving NELF-A.

### SIN3 isoforms share common DNA binding motifs

Neither of the SIN3 isoforms have DNA binding capabilities, but likely are recruited to DNA targets through their association with transcription factors (Kasten et al., 1996). Thus, we conducted a motif analysis to identify enriched motifs at the binding sites of SIN3 isoforms. Furthermore, we parsed out repressed and activated genes to determine if gene regulation outcome is correlated with the binding motif. To do this, we used the MEME software (Buske et al., 2010) through the Galaxy platform. Our analysis found that many binding motifs were different, only 1 out of the top 6 motifs overlapped between the isoforms (Fig. 5, 6). Next, we asked if the SIN3 binding motifs are shared with any transcription factor binding motifs. To this end, we used Tomtom to compare the DNA binding motifs of the SIN3 isoforms with known transcription factor motifs (Gupta et al., 2007). Transcription factors daughterless (Da) and tinman (Tin) binding motifs overlapped with SIN3 220 binding motifs on both activated and repressed genes (Fig. 5A,B). Da is a broadly expressed transcription factor (Cronmiller and Cummings, 1993) with a wide range of regulatory functions, including proliferation (Smith et al., 2002) and development (Cummings and Cronmiller, 1994). *Tin* is a homeobox gene that codes for a transcription factor involved in differentiation and development (Liu et al., 2009; Ranganayakulu et al., 1998; Zaffran et al., 2006). Additionally, the transcription factor longitudinals lacking (Lola) binding motif overlapped with SIN3 187 activated and repressed binding motifs as well as SIN3 220 repressed binding motifs (Fig. 5A, 6). Lola is a transcription factor that plays a role in cell fate determination by antagonizing notch (Zheng and Carthew, 2008). Lola has also been shown to regulate genes involved in programmed cell death (Bass et al., 2007). The finding that multiple transcription factor motifs are enriched at promoters of SIN3-regulated genes are consistent previous results demonstrating that SIN3 recruitment is widespread and that direct gene targets fall into multiple GO categories (Pile and Wassarman, 2000; Saha et al., 2016).

**Fig. 5.**
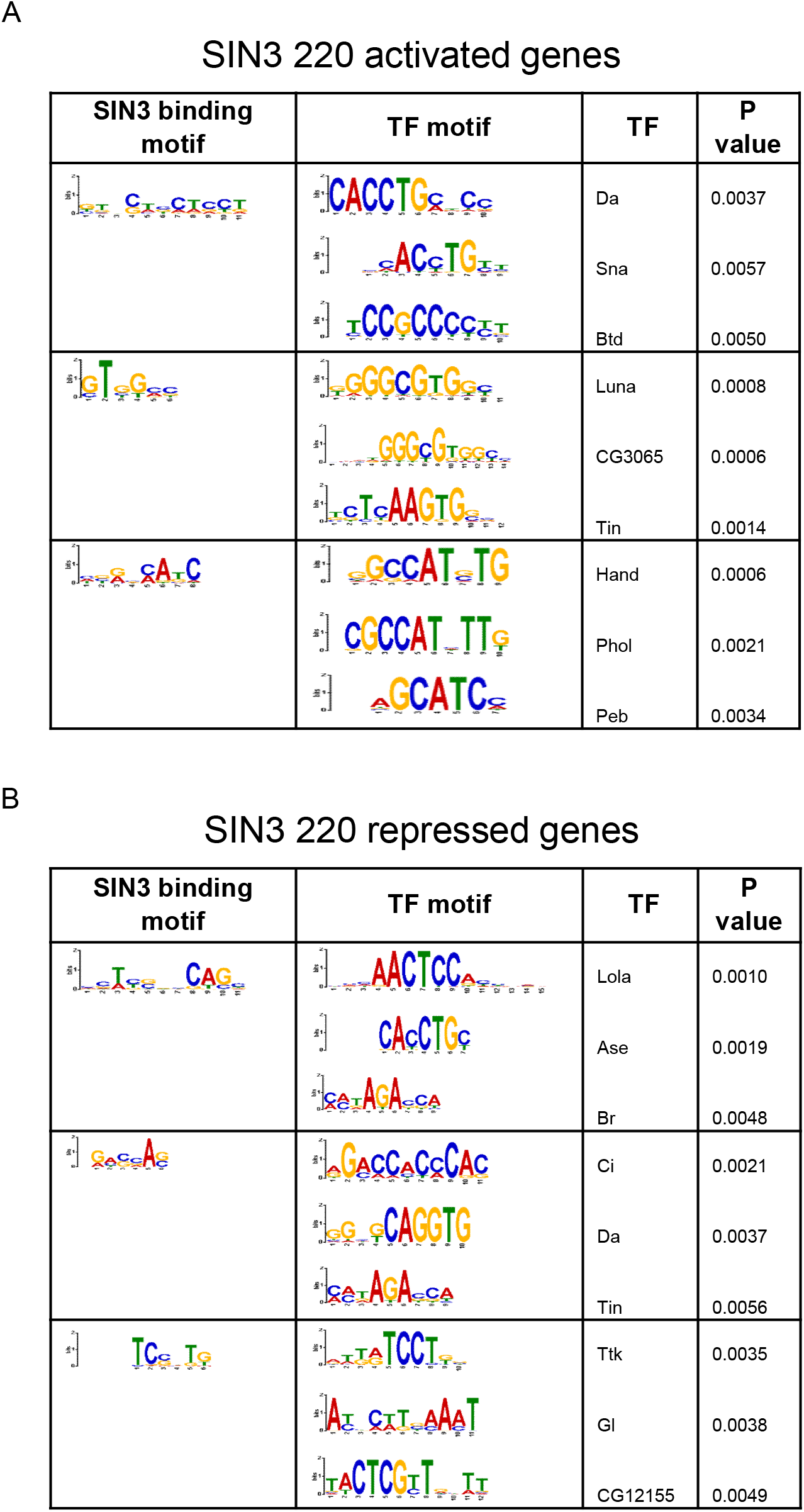
SIN3 220 binding motifs and alignment. A. SIN3 220 activated and B. repressed genes binding motifs were generated using MEME software and top three motifs are shown. Motifs were then compared to known Drosophila transcription factor (TF) motifs using TOMTOM software and the three most statistically significant TFs are shown. Pearson correlation coefficient statistical analysis was utilized (Gupta et al., 2007; Pietrokovski, 1996).

**Fig. 6.**
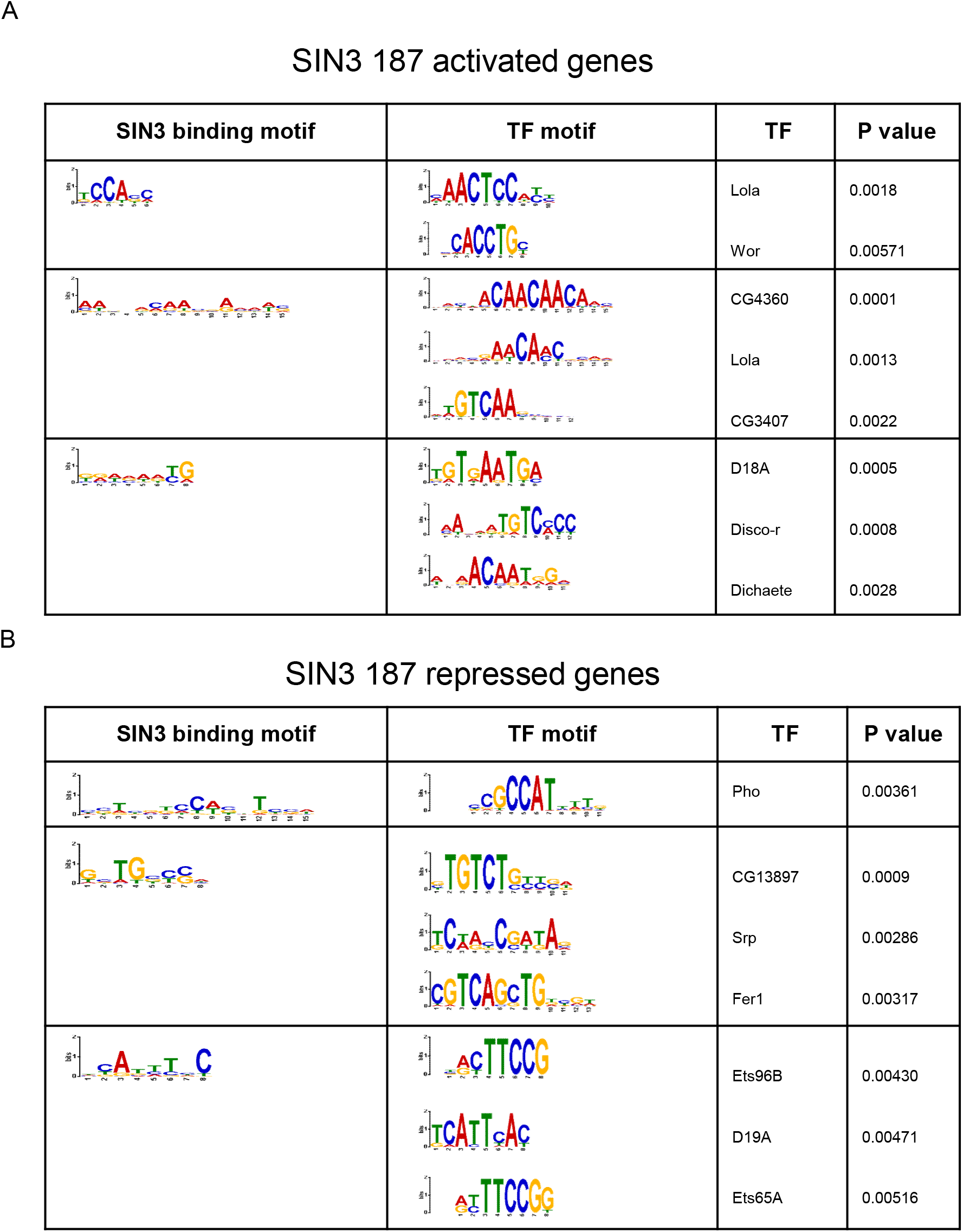
SIN3 187 binding motifs and alignments. Binding motifs of genes A. activated and B. repressed by SIN3 187 were generated using MEME software and top three Motifs are shown. Motifs were then compared to known Drosophila transcription factor (TF) motifs using TOMTOM software and the three most statistically significant TFs are shown. Pearson correlation coefficient statistical analysis was utilized (Gupta et al., 2007; Pietrokovski, 1996).

### SIN3 isoforms function as soft repressors

We previously determined that SIN3 220 functions as a soft repressor (Mitra et al., 2021).This term signifies that SIN3 is capable of fine-tuning the expression of target genes rather than switching them on and off. This action results in small but physiologically and statistically significant changes in gene expression. To expand on our previous study in which we investigated SIN3 220 repressed targets, here we looked at the levels of regulatory action at SIN3 220 activated genes and at genes regulated by SIN3 187. Using direct targets of SIN3 220, we analyzed the log2 fold changes (log2FC) in gene expression of SIN3 220 active and repressed genes. 242 direct repressed targets of SIN3 220 were identified, of which 99.6% (241/242) of genes demonstrated less than 2 log2FC (Fig. 7). A total of 163 directly active targets of SIN3 220 were identified. Of these 163 genes, 98% (160/163) of them showed less than 2 log2FC and only 2% (3/163) of genes fell in the range of 2 to 3 log2FC when levels of SIN3 220 were perturbed (Fig. 7). These data indicate that SIN3 220 indeed acts as a soft regulator on both active as well as repressed gene targets. Next, we asked whether SIN3 187 also functions as a soft regulator to fine-tune gene expression. We hypothesized that since both isoforms have many overlapping targets, they might have similar effects on gene expression. 469 directly repressed targets of SIN3 187 were identified, of which 89.8% (421/469) of genes showed less than 2 log2FC with the overexpression of SIN3 187. 6.4% (30/469) were in the 2 to 3 log2FC category and 3.8% (18/469) of genes showed greater than 3 log2FC in gene expression (Fig. 7). A similar pattern of soft regulatory activity was observed for the direct gene targets activated by SIN3 187. Of the total 390 genes identified, 69.5% (271/390) of genes changed less than 2 log2FC in expression. 13.3% (52/390) showed 2 to 3 log2FC while 17.2% (67/390) of genes had greater than 3 log2FC in expression levels (Fig. 7). These results show that a majority of direct gene targets of both isoforms exhibited small changes in gene expression, demonstrating they function as soft repressors and soft activators. As this mechanism of regulation is seen for repressed as well as activated genes, this modulating activity appears to be the preferred mode of action of this global transcriptional regulator.

**Fig. 7.**
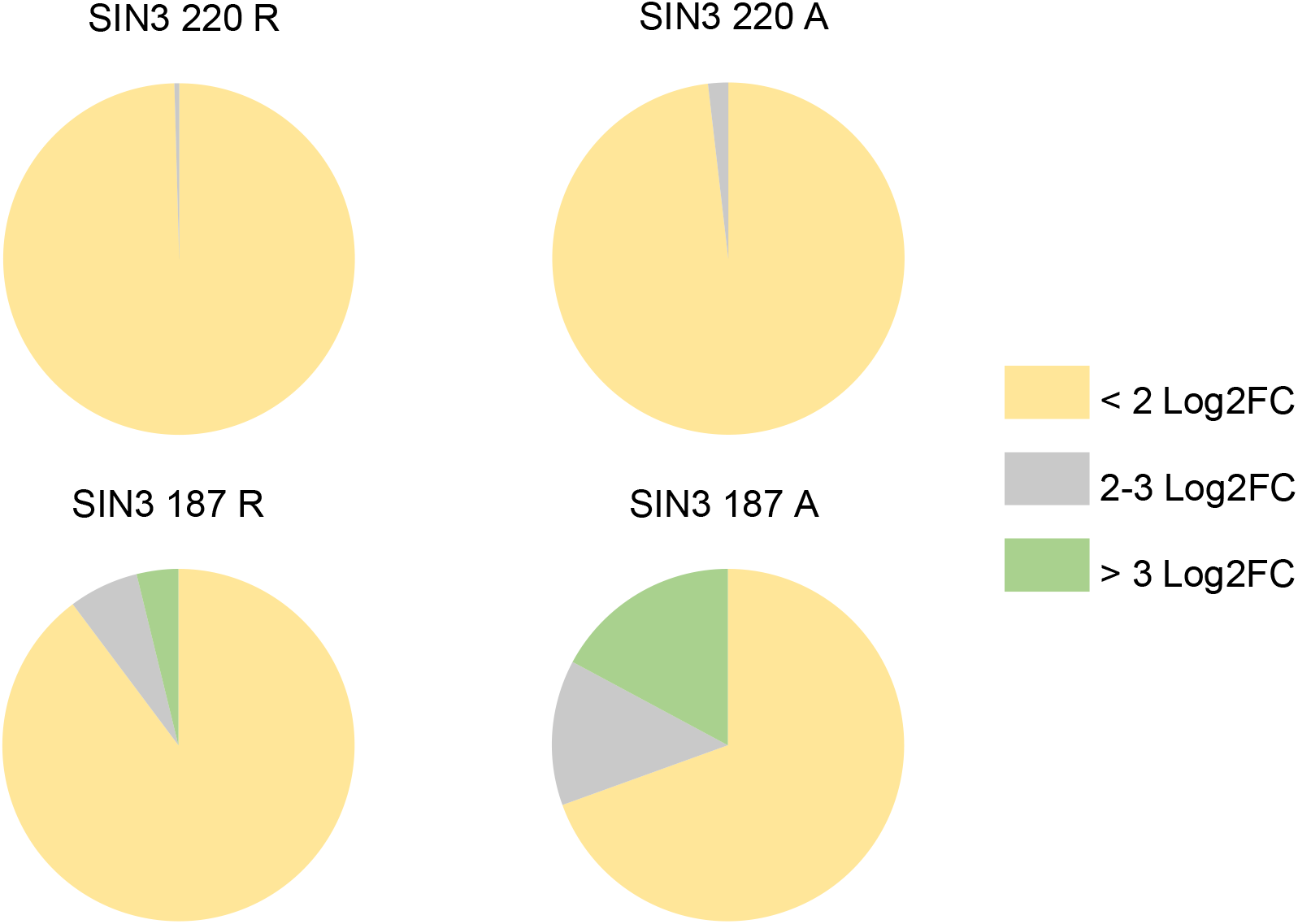
SIN3 isoforms act as soft regulators. Genes regulated and bound by the SIN3 isoforms were parsed based on the expression level change when SIN3 levels were perturbed. R = repressed, A = activated.

### SIN3 fine-tunes the expression of moderately expressed genes

To further investigate whether SIN3 regulates in an on/off or soft regulation manner, we asked if SIN3 regulated gene targets are active or silenced in cells. We hypothesized that if SIN3 is acting as a soft regulator, gene targets should exhibit moderate expression. On the other hand, if SIN3 is acting in an/off manner, gene targets should exhibit either high or silent/low expression, depending on the action of SIN3. To test our hypothesis, we analyzed the expression level of all of the expressed genes in Drosophila S2 cells, as determined by RNA-seq reads reported as fragments per kilobase of exon per million mapped fragments (FPKM). We divided the genes into four quadrants (Fig. 8) using a slightly modified classification as in Kalashnikova and colleagues (Kalashnikova et al., 2021). We considered genes with FPKM >1000 as strongly expressed, genes with FPKM 10-1000 as moderately expressed and genes with FPKM < 1 as silenced (Kalashnikova et al., 2021). We saw that 87% (211/242) of genes repressed by SIN3 220 and 70% (114/163) of genes activated by SIN3 220 were moderately expressed (Fig. 8A). Similarly, to a lesser extent, 87% (406/469) of genes repressed by SIN3 187 and 55% (215/390) of genes activated by SIN3 187 were moderately expressed (Fig. 8A). These data support our hypothesis that SIN3 regulates moderately expressed genes by modulation of activity and not by turning off genes. Interestingly, 14% (53/390) of genes activated following the ectopic expression of SIN3 187 had an FPKM value of 1 or less in control cells, while only 2% of the other gene sets had such a low FPKM value (Fig. 8A). This finding suggests that SIN3 187 may act as a hard regulator on a subset of genes. To further investigate, we looked closely at the 53 genes predicted to be regulated in an on/off manner by SIN3 187. We saw that 96% (51/53) of the genes were activated by more than 3 log2FC following ectopic expression of SIN3 187. Additionally, we asked if these 53 genes activated by SIN3 187 are also regulated by SIN3 220. We found that while 74% (39/53) are bound by SIN3 220, only 6% (3/53) change in expression when SIN3 220 levels are perturbed. These data indicate that SIN3 187 acts as a hard regulator on a subset of genes. To further corroborate our findings, we asked if SIN3 187 binding on those 53 genes is distinct from the binding of SIN3 187 on all SIN3 187 directly activated genes. Comparing the binding of SIN3 187 on the 53 genes with all SIN3 187 directly regulated genes, we saw that SIN3 187 binding on all directly activated genes is localized at the TSS, while SIN3 187 binding on the 53 genes is not strictly localized to the TSS (Fig. 8B). Data presented here provide evidence to suggest that SIN3 187 might have two modes of regulation, soft and hard regulation. We predict that soft regulation occurs due to binding at the promoter of genes, while hard regulation does not.

**Fig. 8.**
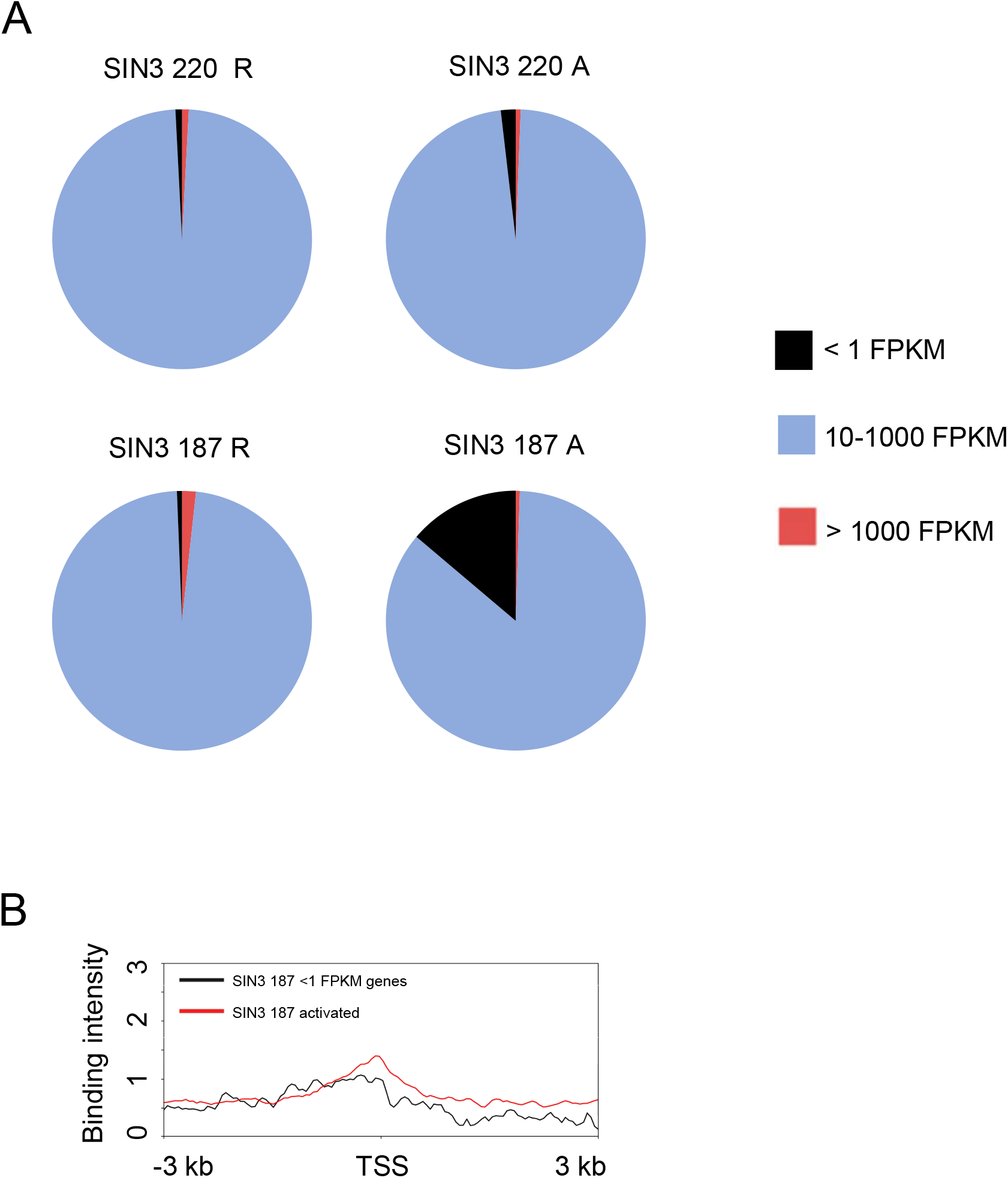
SIN3 187 acts as a hard regulator on a subset of genes. A. FPKM values of genes in Drosophila S2 cells were parsed based on the expression level of those genes in wild-type cells and the regulation of those genes by the SIN3 isoforms. Genes with FPKM >1000 are considered strongly expressed, genes with FPKM 10-1000 as moderately expressed and genes with FPKM < 1 as silenced. B. SIN3 187 binding (from −3 kb to +3 kb from the TSS) on the 53 genes activated by ectopic expression of SIN3 187 and are silent in wild-type S2 cells (black line). This binding profile was compared to the profile of all genes activated by ectopic expression of SIN3 187 (red line). TSS = transcription start site.

## DISCUSSION

Here, we analyzed the binding patterns of the SIN3 isoforms in the context of the chromatin environment. Metagene analysis revealed that the binding of the SIN3 isoforms occurs mostly at the TSS of genes. Interestingly, both SIN3 isoforms exhibited higher binding intensity on genes repressed compared to genes activated (Fig. 2). In contrast, genes that are uniquely repressed by SIN3 187 exhibited a similar binding profile to those genes uniquely activated by SIN3 187. One possible reason for this difference is that SIN3 187 lacks some SIN3 220 complex interactors, some which may help in the recruitment of SIN3 to target genes. Additionally, the difference in binding at the TSS between SIN3 220 and SIN3 187 unique targets leads to the prediction that SIN3 187 can act as a hard regulator while SIN3 220 does not. Soft regulation is predicted to depend on transcriptional regulators binding promoter proximally where they lead to fine-tuning of gene expression (Mitra et al., 2021). Consistent with our hypothesis, soft regulation occurs when SIN3 isoforms are localized to the promoter proximal regions, while hard regulation of genes occurs when SIN3 187 binding is localized away from the TSS. Genes repressed by the SIN3 isoforms exhibited enriched H3K4me3 levels immediately after the TSS (Fig. 3). Given that SIN3 binds at expressed genes, this finding was not surprising since H3K4me3 is enriched on actively expressed genes (Bernstein et al., 2005). The finding that both isoforms are located at targets with similar levels of H3K4me3 enrichment suggests that SIN3 220 and SIN3 187 dampen gene expression through a similar soft regulation mechanism. On the other hand, it is intriguing that both 220 and 187 regulated genes exhibited similar methylation patterns, since the SIN3 220 complex but not the SIN3 187 complex, contains dKDM5/LID (Spain et al., 2010)), a histone demethylase that targets H3K4me3. The H3K4me3 enrichment patterns suggest that H3K4me3 levels at the promoters are not regulated by dKDM5/LID alone.

Furthermore, genes repressed by the SIN3 isoforms exhibited higher RNA pausing at the TSS when compared to genes activated by the isoforms (Fig. 4B). This finding indicates that the mechanism of SIN3 repression is similar between the isoforms and likely involves regulation of RNA Pol II pausing. In line with this prediction, genes repressed by the isoforms showed enrichment of NELF-A, a pausing factor, at the TSS (Fig. 4A). One enzyme that could play a role in the regulation of RNA Pol II binding dynamics is HDAC1. Indeed, both SIN3 isoform complexes contain HDAC1 (Spain et al., 2010). Previously published findings indicate that HDAC activity could lead to RNA Pol II pausing, and the inhibition of HDAC activity leads to the release of the paused polymerase (Vaid et al., 2020). Consistent with the proposed mechanism of regulation, while not statistically significant, SIN3 187 regulated genes had higher average RNA Pol II pausing when compared to SIN3 220 (Fig. 4B). We previously determined that the Vmax of the HDAC activity of the SIN3 187 complex is higher when compared to the SIN3 220 complex (Spain et al., 2010). We predict that the higher HDAC activity in the SIN3 187 complex is responsible for a higher pausing index when compared to the SIN3 220 complex. Overall, data presented here along with published data, allow us to propose a model in which SIN3 repression activity is in part dependent on the role of HDAC1 in regulating RNA Pol II pausing.

Soft regulation is predicted to aftect gene expression in a less dramatic but biologically significant manner. Our findings support the role of both SIN3 220 and SIN3 187 in soft regulation. The perturbation of either isoform leads to small but significant changes in the level of expression of target genes (Fig. 7). Furthermore, both isoforms predominantly bind at the TSS of genes, a feature predicted of soft regulation (Mitra et al., 2021). Through analysis of the measured expression level of genes regulated by the SIN3 isoforms, we observed that most of the gene targets are expressed at moderate levels in S2 cells. This finding is consistent with our idea of soft regulation, wherein genes expressed at moderate levels are attenuated and not turned on/off. Interestingly, a subset of genes activated by SIN3 187 exhibited the opposite effect (Fig. 8B). These genes were not expressed in wild-type cells and are turned on by SIN3 187 to more than 3 log2 fold change. These data indicate that the SIN3 187 complex might regulate genes by two mechanisms, soft and hard regulation. Future studies will be aimed at further testing the model that the SIN3 complexes impact histone modifications at housekeeping gene targets to modulate expression levels in response to cellular demands.

## MATERIALS AND METHODS

All of the software packages were used through the public servers at the Galaxy web platform (Afgan et al., 2018)) with the exception of the RNA Pol II pausing index calculation and Tomtom.

### ChlP-Seq analysis

Data were downloaded from the Sequence Read Archive (SRA), using NCBI SRA toolkit (Leinonen et al., 2011). To determine the binding profiles, we first mapped all reads to the *Drosophila melanogaster* reference genome (dm3) using the software package Burrows-Wheeler Aligner (BWA) (Li and Durbin, 2009). Uniquely mapped reads were extracted using the filter SAM option in SAMtools (Li et al., 2009).

### Heat maps and binding profiles

Using the UCSC table browser (Karolchik et al., 2004), we created a genome assembly using a list of genes of interest such as SIN3 220 activated genes, SIN3 220 repressed genes, and so on. ChlP-seq data was then mapped to the assembled genome and a matrix was created using deepTools2: computeMatrix (Ramirez et al., 2016). This matrix, centered around the TSS and +/- 3 kb, was then used to generate heat maps and graphs plotting the binding profiles of the protein or histone mark of interest.

### Pausing index

Rpb3 enrichment on SIN3-regulated genes was calculated and placed into a matrix using deepTools2: computeMatrix (Ramirez et al., 2016), through the Galaxy web platform. This matrix was then downloaded from Galaxy and analyzed through R Studio. Rpb3 enrichment was then calculated at the promoter region (−50 bp to +50 bp) and divided by the genic region (+300 bp to −100 bp TES). Statistical analysis using the Mann-Whitney test was done.

### MEME and Tomtom analysis

MACS2 (version 2.1.1) (Zhang et al., 2008) was used to call peaks using the default parameters. Genomic locations were extracted and binding motifs were determined using MEME suite (Bailey and Elkan, 1994). To compare the binding motifs of SIN3 isoforms to transcription factor binding motifs, Tomtom package using default settings was used (Tanaka et al., 2011).

### FPKM levels of genes in S2 cells

RNA-seq data from control cells previously published by our group (Gajan et al., 2016) was downloaded and divided into five categories based on the FPKM as previously published (Kalashnikova et al., 2021).

## ACKNOWLEDGEMENTS

We thank Dr. Roger Pique-Regi, Wayne State University, for his assistance in calculating the pausing index. We are indebted to Dr. Laure Teysset (Sorbonne Université) for teaching us how to utilize the Galaxy platform and to Dr. Olivier Lespinet and Dr. Melina Gallopin (Université Paris-Saclay) for introducing us to Rstudio for analysis. We thank all members of the Pile laboratory for their evaluation of this manuscript.

## AUTHOR CONTRIBUTIONS

I.S and L.A.P conceived the project. I.S and A.M performed the analysis. I.S and L.A.P wrote and edited the manuscript.

## DATA AVAILABILITY

All data used in the study are available through the GEO database. GEO accession numbers are listed in Figure 1A.

